# Functional screening of β-Glucanase Producing Actinomycetes Strains from Western Ghats ecosystems of Kerala, India

**DOI:** 10.1101/2020.04.11.036731

**Authors:** Lekshmi K. Edison, N. S. Pradeep

## Abstract

Screening of potential soil actinomycetes is static at infant phase because less than one part of soil biodiversity has been explored. An important factor considered before isolating microorganisms with potential application is understanding the biodiversity and environmental features associated with growth. Search of distinctive enzymes from unusual ecological habitats are highly fascinating and have great opportunities that may also pointed the developments in high throughput screening programs. In the present study Western Ghats hot spot regions of Kerala has been explored for the actinomycetes strains with beta glucanase activity. A total of 127 actinomycetes strains were isolated. After qualitative primary screening 106 strains (83%) produced exo-β-1,4-glucanase enzyme and 79 strains (62%) produced endo-β-1,3-glucanase enzyme. The quantitative secondary screening confirmed the strains TBG-MR17 and TBG-AL13 recognised as respective dominant producers of exo-β-1,4-glucanase and endo-β-1,3-glucanase enzymes. The study reveals the richness of the Western Ghats soils with innumerable actinomycetes having potential β-glucanase activities.

## Introduction

Western Ghats in India are well-known biodiversity hot spot of rich flora and fauna, also recognised highly productive ecosystems. It is a forested strip of relatively old mountain ranges, beginning from Central Maharashtra and stretched up to the Southern tip of Kerala. These areas are granted with a “heritage tag” by UNESCO as a gene pool, sheltering millions of species of animals, plants and microbes. Western Ghats regions are domicile to immense collection of unexplored and novel microbial diversity including actinomycetes species. Exploitation of unique, natural, highly documented and less explored biodiversity ecosystems for actinomycetes is highly necessary for the discovery of novel bioactive metabolites with prospective applications (Jalaja et al., 2011; Mohandas et al., 2012; Balachandran et al., 2012; Nampoothiri et al., 2013).

Microbial β-glucanase have been isolated from variety of microbes and well characterized (Velho-Pereira and Kamat, 2013). Actinomycetes have been extensively recognised as a source of β-glucan degrading enzymes. Among the wide genus, Streptomyces are most prevalent group of enzyme producers (Wu et al., 2018). Some of them includes Streptomyces sioyaensis (Hong et al., 2008), Streptomyces sp. 9×166 (Sakdapetsiri et al., 2016), Streptomyces sp. S27 (Shi et al., 2010), Streptomyces matensis ATCC 23935 (Woo et al., 2014), Streptomyces rochei (Wu et al., 2002) and Streptomyces sp. EF-14 (Fayad et al., 2001).

Nevertheless β-glucanase are less characterized in actinomycetes strains within the Western Ghats regions. These regions remain less explored, and also due to inextricable altered habitat the chances of obtaining potential strains of actinomycetes including Streptomyces species with exceptional β-glucanase activities are much higher. This chapter deals with the isolation of actinomycetes strains by exploring selected Western Ghats regions of Kerala for quantitative and qualitative screening of two β-glucanase enzymes, exo-β-1,4-glucanase and endo-β-1,3-glucanase. The synergic action of both β-glucanases requires the complete degradation of barley and oat β-glucans, which is essential for industrial applications such as brewing industry and feed enzyme industry.

## Materials and Methods

### Sample Collection

Soil samples were collected from Western Ghats of Kerala; include Wayandu, Munnar, Chinnar, Marayoor, Anamala, Neryamangalam, Nelliyampathi, Agasthyarkoodum, Palode and Kulathuppuzha. After removing the surface layer (approximately top 4 cm), the soil samples were taken from a depth of 5 to 10 cm of the superficial layers in each location. Three different samples were collected from each areas.

### Pre-treatment of Soil Samples and Isolation of Actinomycetes Strains

Soil samples were pre-treated with 1% CaCO_3_ incubated at 28°C for 10 days before use (El-Nakeeb and Lechevalier, 1963). One gram of soil taken in 100mL Erlenmeyer flask was pre-treated by heating the soil at 50°C for 1 h. Standard serial dilution plate method was employed for the isolation of actinomycetes strains. 9mL sterile distilled water was added to the 1g of previously oven dried soil samples and mixed thoroughly in a rotary shaker for 30 min at 150 rpm at room temperature. The suspension was serially diluted to obtain 10-3 to 10-7 dilutions. 1.0mL of each dilution was pour plated on starch casein agar (SCA) plates in triplicate. After incubation at 28°C for 7 days the actinomycetes colonies were counted and represented as colony-forming units per gram (CFU.g-1) of soil. For the purification of single isolated colonies, streak plate method was used.

### Primary Screening for Exo-β-1,4-Glucanase and Endo-β-1,3-Glucanase Activities

Plate assay method was used for primary screening of enzymes (Meddeb-Mouelhi et al., 2014). Isolated actinomycetes strains were spot inoculated on a modified agar screening media with 1% (w/v) Avicel® PH-101 (Sigma) as a substrate for exo-β-1,4-glucanase and 0.2% (w/v) AZCL-Pachyman (Megazyme, Germany) as substrate for endo-β-1,3-glucanase enzymes and incubated for 5 days at 28°C. The presence of clear zone around the growth indicated the exo-β-1,4-glucanase activity. The enzymatic index (EI) of each strain was calculated as follows by measuring the diameter of hydrolysis zone and diameter of colony.

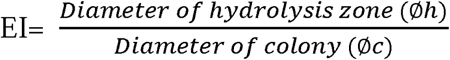

The experiment was performed using three replicates for each strain. The average EI of three experiments were calculated, together with standard deviation.

### Secondary Screening for Exo-β-1,4-Glucanase and Endo-β-1,3-Glucanase Activities

Modified liquid media with 0.5% (w/v) Avicel (for exo-β-1,4-Glucanase) and 0.2% (w/v) CM-curdlan (for endo-β-1,3-Glucanase) were used. The enzyme activities were quantitatively estimated by DNS assay method by measuring the released reducing sugars (Miller, 1959). One unit of beta glucanase activity is defined as the amount of enzyme required to release 1 µmol reducing sugar (as glucose equivalence) in one minute under defined conditions (Fulop and Ponyi, 1997).

## Results

### Isolation of Actinomycetes Strains

The study explored Western Ghats biodiversity for the isolation of β-glucanase producing actinomycetes strains. 10 different areas of Western Ghats of Kerala such as Wayanad, Nelliyampathy, Neriyamangalam, Munnar, Chinnar, Anamalai, Marayoor, Kulathupuzha, Palode and Agasthyarkoodam, were explored. All the selected sample collection spots were mountain and natural forest hot spot areas. A total of 127 morphologically different actinomycetes strains were isolated. The isolates formed well branched substrate mycelia and ample aerial hyphae that segregated into well-developed spores chains. The number of actinomycetes strains obtained per gram of soil samples from these areas are shown in Table1.

**Table 1.** Description of soil sampling areas and number of colonies

| Area | Description | Actinomycetes Colonies (CFU.g<sup>-1</sup>) |
| --- | --- | --- |
| Wayanad | Mountain forest | 15x10<sup>-3</sup> |
| Nelliyampathy | Hill range | 10x10<sup>-3</sup> |
| Neryamangalam | Natural forest | 7x10<sup>-3</sup> |
| Munnar | Hill station | 11x10<sup>-3</sup> |
| Chinnar | Lower mountain forest | 23x10<sup>-3</sup> |
| Anaimalai | Mountain | 24x10<sup>-3</sup> |
| Marayur | Natural sandal wood forest | 5x10<sup>-3</sup> |
| Kulathupuzha | Natural forest | 13x10<sup>-3</sup> |
| Palode | Reserve forest | 5x10<sup>-3</sup> |
| Agasthyakoodam | Hill range | 14x10<sup>-3</sup> |

### Primary Screening of Exo-β-1,4-Glucanase and Endo-β-1,3-Glucanase Activities

All the isolated actinomycetes strains were evaluated for semi-quantitative production of exo-β-1,4-glucanase and endo-β-1,3-glucanase activities using plate assay method. Exo-β-1,4-glucanase production was screened using 1% Avicel (microcrystalline cellulose) as sole carbon source. The presence of a pale halo around the colonies after congo red dye staining indicated the production of enzyme exo-β-1,4-glucanase. This zone of clearance was due hydrolysis of Avicel by exo-β-1,4-glucanase to glucose residues, it does not have any affinity to congo red. Out of 127 isolates, 106 strains (83% of total strains) produced exo-β-1,4-glucanase enzyme activity (Table 2).

**Table 2.** Primary screening of exo-β-1,4-glucanase activity

| No | Strain | Mean<br>Øh | Mean Øc | Mean EI | SD |
| --- | --- | --- | --- | --- | --- |
| 1 | TBG-CH1 | 2.9 | 0.7 | 4.10 | 0.089 |
| 2 | TBG-CH2 | 2.4 | 0.7 | 3.40 | 0.089 |
| 3 | TBG-CH3 | 3 | 0.7 | 4.30 | 0.155 |
| 4 | TBG-CH4 | 3 | 0.9 | 3.30 | 0.179 |
| 5 | TBG-CH5 | 3.3 | 0.9 | 3.73 | 0.137 |
| 6 | TBG-CH6 | 3.2 | 1 | 3.23 | 0.186 |
| 7 | TBG-CH7 | 3.3 | 0.8 | 4.13 | 0.137 |
| 8 | TBG-CH8 | 3.4 | 0.9 | 3.80 | 0.089 |
| 9 | TBG-CH9 | No zone |  |  |  |
| 10 | TBG-CH10 | No zone |  |  |  |
| 11 | TBG-CH11 | 3 | 0.9 | 3.33 | 0.258 |
| 12 | TBG-CH12 | 2.4 | 0.7 | 3.40 | 0.089 |
| 13 | TBG-CH13 | 2.3 | 1 | 2.30 | 0.089 |
| 14 | TBG-CH14 | 2 | 0.6 | 3.30 | 0.089 |
| 15 | TBG-CH15 | 2.9 | 0.7 | 4.13 | 0.052 |
| 16 | TBG-CH16 | 2.5 | 0.6 | 4.20 | 0.089 |
| 17 | TBG-CH17 | 3.3 | 0.9 | 3.63 | 0.052 |
| 18 | TBG-CH18 | 3.1 | 0.8 | 4.10 | 0.322 |
| 19 | TBG-CH19 | 1.8 | 0.8 | 2.23 | 0.052 |
| 20 | TBG-CH20 | 3.1 | 0.8 | 4.00 | 0.089 |
| 21 | TBG-CH21 | 2.5 | 0.6 | 4.20 | 0.179 |
| 22 | **TBG-CH22\*** | **2.7** | **0.6** | **4.50** | **0.089** |
| 23 | TBG-CH23 | 3.2 | 0.8 | 4.00 | 0.089 |
| 24 | TBG-AL1 | 1.5 | 0.9 | 1.73 | 0.052 |
| 25 | TBG-AL2 | No zone |  |  |  |
| 26 | TBG-AL3 | 2.3 | 1.2 | 1.87 | 0.052 |
| 27 | TBG-AL4 | 1.5 | 0.8 | 1.90 | 0.179 |
| 28 | TBG-AL5 | 2.9 | 1.2 | 2.40 | 0.155 |
| 29 | TBG-AL6 | 3 | 0.9 | 3.30 | 0.089 |
| 30 | TBG-AL7 | 3.5 | 1.2 | 2.90 | 0.089 |
| 31 | TBG-AL8 | 1.75 | 0.7 | 2.50 | 0.089 |
| 32 | TBG-AL9 | 2.8 | 0.7 | 4.03 | 0.052 |
| 33 | TBG-AL10 | 3.15 | 0.8 | 3.87 | 0.052 |
| 34 | TBG-AL11 | 2.1 | 0.8 | 2.60 | 0.089 |
| 35 | TBG-AL12 | 3 | 0.9 | 3.30 | 0.089 |
| 36 | TBG-AL13 | 3.5 | 1.3 | 2.73 | 0.137 |
| 37 | TBG-AL14 | 1.8 | 0.8 | 2.30 | 0.155 |
| 38 | TBG-AL16 | 2.5 | 1.2 | 2.10 | 0.155 |
| 39 | TBG-AL17 | 1.5 | 0.5 | 3.03 | 0.052 |
| 40 | TBG-AL18 | 2.7 | 0.7 | 3.93 | 0.186 |
| 41 | **TBG-AL19\*** | **3.6** | **0.6** | **6.03** | **0.137** |
| 42 | TBG-AL20 | 2.6 | 0.7 | 3.67 | 0.052 |
| 43 | TBG-AL21 | 2.8 | 1 | 2.80 | 0.089 |
| 44 | TBG-AL22 | 2.5 | 1 | 2.50 | 0.089 |
| 45 | TBG-AL23 | 2.5 | 0.9 | 2.83 | 0.052 |
| 46 | TBG-AL24 | 3.1 | 0.9 | 3.40 | 0.089 |
| 47 | TBG-AL25 | 2.1 | 0.8 | 2.60 | 0.089 |
| 48 | **TBG-NR1\*** | **3.7** | **0.8** | **4.63** | **0.052** |
| 49 | **TBG-NR2\*** | **3.7** | **0.8** | **4.63** | **0.137** |
| 50 | **TBG-NR3\*** | **2.9** | **0.4** | **7.27** | **0.225** |
| 51 | TBG-NR4 | 1.7 | 0.5 | 3.37 | 0.137 |
| 52 | TBG-NR6 | 3.9 | 1 | 3.90 | 0.089 |
| 53 | TBG-NR7 | 3 | 1 | 2.97 | 0.186 |
| 54 | TBG-NR11 | 3.6 | 0.9 | 4.00 | 0.179 |
| 55 | TBG-NR14 | 3.6 | 0.9 | 3.90 | 0.089 |
| 56 | TBG-NR16 | 2.8 | 0.9 | 3.07 | 0.052 |
| 57 | TBG-NR18 | 3.7 | 0.9 | 4.07 | 0.103 |
| 58 | **TBG-NR19\*** | **3.6** | **0.8** | **4.47** | **0.052** |
| 59 | **TBG-NR21\*** | **3.8** | **0.8** | **4.80** | **0.179** |
| 60 | TBG-NR22 | No zone |  |  |  |
| 61 | **TBG-NR23\*** | **3.6** | **0.7** | **5.10** | **0.089** |
| 62 | **TBG-NR24\*** | **3.6** | **0.8** | **4.50** | **0.089** |
| 63 | TBG-MN1 | 3.4 | 0.9 | 3.83 | 0.137 |
| 64 | TBG-MN2 | 3.3 | 0.8 | 4.07 | 0.207 |
| 65 | TBG-MN3 | 3.7 | 1.2 | 3.13 | 0.137 |
| 66 | **TBG-MN5\*** | **3.3** | **0.7** | **4.67** | **0.186** |
| 67 | TBG-MN6 | 3.5 | 1 | 3.47 | 0.137 |
| 68 | TBG-MN7 | 3 | 0.8 | 3.80 | 0.089 |
| 69 | TBG-MN8 | 3.3 | 0.8 | 4.13 | 0.137 |
| 70 | TBG-MN9 | 3.4 | 0.8 | 4.27 | 0.137 |
| 71 | TBG-MN10 | 3.1 | 0.9 | 3.40 | 0.089 |
| 72 | TBG-MN11 | 3.5 | 1 | 3.53 | 0.137 |
| 73 | TBG-MN12 | 3.3 | 0.8 | 4.13 | 0.137 |
| 74 | TBG-MR1 | 3.7 | 0.9 | 4.07 | 0.137 |
| 75 | TBG-MR2 | 2.9 | 0.8 | 3.57 | 0.052 |
| 76 | **TBG-MR3\*** | **3.8** | **0.8** | **4.77** | **0.137** |
| 77 | TBG-MR4 | 3.5 | 1 | 3.47 | 0.052 |
| 78 | TBG-MR5 | 2.8 | 1 | 2.77 | 0.052 |
| 79 | TBG-MR7 | 2.2 | 1 | 2.17 | 0.137 |
| 80 | **TBG-MR8\*** | **4** | **0.8** | **5.03** | **0.052** |
| 81 | **TBG-MR9\*** | **4.2** | **0.9** | **4.67** | **0.186** |
| 82 | TBG-MR12 | 3.5 | 1 | 3.50 | 0.089 |
| 83 | **TBG-MR17\*** | **3.5** | **0.7** | **5.03** | **0.137** |
| 84 | TBG-MR18 | 2.8 | 0.8 | 3.47 | 0.052 |
| 85 | TBG-MR19 | 3 | 0.8 | 3.75 | 0.089 |
| 86 | TBG-I5II | 3.7 | 0.9 | 4.10 | 0.089 |
| 87 | TBG-N1 | 3.3 | 0.8 | 4.12 | 0.207 |
| 88 | TBG-N2 | 3.4 | 0.8 | 4.20 | 0.155 |
| 89 | TBG-N3 | No zone |  |  |  |
| 90 | TBG-N5 | No zone |  |  |  |
| 91 | TBG-N6 | 3.6 | 0.8 | 4.33 | 0.273 |
| 92 | TBG-N7 | No zone |  |  |  |
| 93 | TBG-N8 | 2.7 | 0.8 | 3.40 | 0.155 |
| 94 | TBG-NMP6 | No zone |  |  |  |
| 95 | TBG-NMP5 | 3.4 | 1.2 | 2.80 | 0.089 |
| 96 | TBG-NMP7 | No zone |  |  |  |
| 97 | TBG-AM1A | 2.5 | 0.9 | 2.80 | 0.089 |
| 98 | TBG-AM5 | 3.5 | 0.8 | 4.40 | 0.089 |
| 99 | TBG-AM31 | 3.7 | 1.1 | 3.40 | 0.155 |
| 100 | TBG-AM21 | 2.4 | 0.8 | 3.00 | 0.390 |
| 101 | TBG-AM27 | No zone |  |  |  |
| 102 | TBG201 | 2.8 | 0.8 | 3.50 | 0.390 |
| 103 | TBG19 | No zone |  |  |  |
| 104 | TBG3-6 | 3.4 | 1.2 | 2.80 | 0.179 |
| 105 | TBG3-8 | No zone |  |  |  |
| 106 | TBG3-12 | No zone |  |  |  |
| 107 | TBG-AM47 | No zone |  |  |  |
| 108 | TBG-AM83 | 3.3 | 0.8 | 4.10 | 0.473 |
| 109 | TBG-AM2 | 2.3 | 0.4 | 3.80 | 0.237 |
| 110 | TBG-AM42 | No zone |  |  |  |
| 111 | TBG-AM55 | 3.5 | 1 | 3.50 | 0.179 |
| 112 | TBG-AM57 | No zone |  |  |  |
| 113 | TBG-AG21 | 3.7 | 0.6 | 4.03 | 0.186 |
| 114 | TBG-AG20 | No zone |  |  |  |
| 115 | TBG-AG28 | 3.7 | 0.9 | 4.10 | 0.089 |
| 116 | TBG-B1 | No zone |  |  |  |
| 117 | TBG-B2 | 2.3 | 1.3 | 1.80 | 0.237 |
| 118 | TBG-B3 | No zone |  |  |  |
| 119 | TBG-B4 | No zone |  |  |  |
| 120 | TBG-B5 | No zone |  |  |  |
| 121 | TBG-WY1 | No zone |  |  |  |
| 122 | TBG-WY2 | 3.1 | 0.9 | 3.40 | 0.358 |
| 123 | TBG-WY5 | 3.1 | 0.9 | 3.40 | 0.358 |
| 124 | TBG-WY8 | 2.7 | 0.8 | 3.40 | 0.358 |
| 125 | TBG-WY9 | 2.7 | 0.6 | 4.50 | 0.322 |
| 126 | TBG-WY17 | 2.5 | 0.9 | 2.80 | 0.237 |
| 127 | TBG-WY19 | No zone |  |  |  |

\*strains selected for secondary screening

The endo-β-1,3-glucanase activity of isolated actinomycetes strains were screened using 0.2% AZCL-Pachyman as a carbon source in plate assay. AZCL-Pachyman is an insoluble blue coloured azurine dye cross-linked substrate specifically for the determination of endo-β-1,3-glucanase activity. The endo-β-1,3-glucananolytic activity of actinomycetes strains were produced a blue halo around the colonies indicating the zone of hydrolysis. This is due to the activity of enzyme on insoluble AZCL-Pachyman released water soluble blue colour dyed fragments. Among the total 127 isolated strains, only 79 strains (62%) showed endo-β-1,3-glucanase activity (Table 3). The EI value of exo-β-1,4-glucanase was shown in between 1.7 to 7.3 and that of endo-β-1,3-glucanase was in between 1.8 to 8.5. Strains showed high EI values (in and above 4.5) were considered as potential enzyme producers and were selected for secondary screening (quantitative screening).

**Table 3.** Primary screening of endo-β-1,3-glucanase activity

| No | Strain | Mean Øh | Mean Øc | Mean EI | SD |
| --- | --- | --- | --- | --- | --- |
| 1 | TBG-CH1 | 2.2 | 0.6 | 3.60 | 0.200 |
| 2 | TBG-CH2 | No zone |  |  |  |
| 3 | TBG-CH3 | 2 | 0.8 | 2.50 | 0.300 |
| 4 | TBG-CH4 | No zone |  |  |  |
| 5 | TBG-CH5 | 1.9 | 0.9 | 2.13 | 0.252 |
| 6 | TBG-CH6 | 2.2 | 0.6 | 3.60 | 0.300 |
| 7 | TBG-CH7 | 1.5 | 0.5 | 3.00 | 0.300 |
| 8 | TBG-CH8 | No zone |  |  |  |
| 9 | TBG-CH9 | 2.4 | 1 | 2.43 | 0.058 |
| 10 | TBG-CH10 | No zone |  |  |  |
| 11 | TBG-CH11 | No zone |  |  |  |
| 12 | TBG-CH12 | 1.4 | 0.7 | 2.03 | 0.058 |
| 13 | TBG-CH13 | No zone |  |  |  |
| 14 | TBG-CH14 | No zone |  |  |  |
| 15 | TBG-CH15 | No zone |  |  |  |
| 16 | TBG-CH16 | No zone |  |  |  |
| 17 | TBG-CH17 | No zone |  |  |  |
| 18 | TBG-CH18 | No zone |  |  |  |
| 19 | TBG-CH19 | No zone |  |  |  |
| 20 | TBG-CH20 | 2.2 | 0.8 | 2.80 | 0.100 |
| 21 | TBG-CH21 | 2.4 | 0.7 | 3.40 | 0.458 |
| 22 | **TBG-CH22\*** | **1.7** | **0.2** | **8.53** | **0.058** |
| 23 | TBG-CH23 | 1.8 | 0.6 | 3.00 | 0.346 |
| 24 | TBG-AL1 | 2.2 | 0.7 | 3.10 | 0.265 |
| 25 | TBG-AL2 | 2.5 | 0.8 | 3.10 | 0.346 |
| 26 | TBG-AL3 | 2.4 | 0.9 | 2.70 | 0.265 |
| 27 | TBG-AL4 | 3.2 | 1.1 | 2.90 | 0.200 |
| 28 | TBG-AL5 | 1.8 | 0.7 | 2.60 | 0.361 |
| 29 | TBG-AL6 | No zone |  |  |  |
| 30 | TBG-AL7 | No zone |  |  |  |
| 31 | TBG-AL8 | 1.8 | 0.5 | 3.60 | 0.173 |
| 32 | TBG-AL9 | No zone |  |  |  |
| 33 | TBG-AL10 | No zone |  |  |  |
| 34 | TBG-AL11 | No zone |  |  |  |
| 35 | TBG-AL12 | 2.3 | 0.8 | 2.90 | 0.200 |
| 36 | **TBG-AL13\*** | **2.6** | **0.4** | **6.90** | **0.100** |
| 37 | **TBG-AL14\*** | **3.4** | **0.8** | **4.60** | **0.458** |
| 38 | TBG-AL16 | No zone |  |  |  |
| 39 | TBG-AL17 | No zone |  |  |  |
| 40 | TBG-AL18 | No zone |  |  |  |
| 41 | TBG-AL19 | 2.8 | 0.9 | 3.10 | 0.265 |
| 42 | TBG-AL20 | 2.2 | 0.8 | 2.80 | 0.200 |
| 43 | TBG-AL21 | No zone |  |  |  |
| 44 | TBG-AL22 | 2 | 1 | 2.00 | 0.300 |
| 45 | TBG-AL23 | 2.5 | 0.7 | 3.90 | 0.265 |
| 46 | TBG-AL24 | 2.5 | 0.6 | 4.10 | 0.173 |
| 47 | TBG-AL25 | 2.3 | 0.8 | 2.80 | 0.200 |
| 48 | TBG-NR1 | 3.2 | 0.9 | 3.50 | 0.361 |
| 49 | **TBG-NR2\*** | **2.3** | **0.5** | **4.60** | **0.346** |
| 50 | TBG-NR3 | No zone |  |  |  |
| 51 | TBG-NR4 | No zone |  |  |  |
| 52 | TBG-NR6 | 3 | 0.9 | 3.30 | 0.300 |
| 53 | TBG-NR7 | No zone |  |  |  |
| 54 | TBG-NR11 | 3.5 | 0.8 | 4.30 | 0.200 |
| 55 | **TBG-NR14\*** | **1.5** | **0.3** | **5.00** | **0.600** |
| 56 | TBG-NR16 | No zone |  |  |  |
| 57 | TBG-NR18 | 3 | 0.9 | 3.30 | 0.265 |
| 58 | TBG-NR19 | 2.8 | 1 | 2.80 | 0.100 |
| 59 | TBG-NR21 | 3.1 | 0.9 | 3.40 | 0.361 |
| 60 | TBG-NR22 | No zone |  |  |  |
| 61 | TBG-NR23 | 3.6 | 1.1 | 3.20 | 0.361 |
| 62 | **TBG-NR24\*** | **2.5** | **0.5** | **5.00** | **0.600** |
| 63 | TBG-MN1 | 2.2 | 0.7 | 3.10 | 0.200 |
| 64 | TBG-MN2 | 1.9 | 0.9 | 2.10 | 0.200 |
| 65 | TBG-MN3 | 3.5 | 0.9 | 3.80 | 0.265 |
| 66 | TBG-MN5 | 2.7 | 0.7 | 3.80 | 0.265 |
| 67 | TBG-MN6 | 2.6 | 1 | 2.60 | 0.300 |
| 68 | TBG-MN7 | 1.9 | 0.5 | 3.80 | 0.200 |
| 69 | TBG-MN8 | 2.5 | 0.6 | 4.10 | 0.361 |
| 70 | TBG-MN9 | 2.2 | 0.8 | 2.75 | 0.466 |
| 71 | TBG-MN10 | 2.5 | 0.8 | 3.10 | 0.265 |
| 72 | TBG-MN11 | 2.5 | 1 | 2.50 | 0.265 |
| 73 | TBG-MN12 | No zone |  |  |  |
| 74 | TBG-MR1 | 2 | 0.8 | 2.57 | 0.306 |
| 75 | **TBG-MR2\*** | **2.3** | **0.5** | **4.60** | **0.100** |
| 76 | TBG-MR3 | 2.5 | 0.8 | 3.10 | 0.100 |
| 77 | TBG-MR4 | 2.1 | 0.7 | 3.00 | 0.173 |
| 78 | TBG-MR5 | 1.3 | 0.6 | 2.10 | 0.100 |
| 79 | **TBG-MR7\*** | **2.4** | **0.5** | **4.80** | **0.300** |
| 80 | TBG-MR8 | 2.9 | 1 | 2.90 | 0.200 |
| 81 | TBG-MR9 | No zone |  |  |  |
| 82 | TBG-MR12 | 2.6 | 0.8 | 3.20 | 0.400 |
| 83 | TBG-MR17 | 2.8 | 0.9 | 3.10 | 0.265 |
| 84 | TBG-MR18 | No zone |  |  |  |
| 85 | TBG-MR19 | 1.5 | 0.6 | 2.50 | 0.265 |
| 86 | TBG-I5II | 1.8 | 0.7 | 2.60 | 0.200 |
| 87 | TBG-N1 | 2.2 | 0.7 | 3.10 | 0.100 |
| 88 | TBG-N2 | 2.1 | 0.6 | 3.50 | 0.300 |
| 89 | TBG-N3 | 2.7 | 0.7 | 3.90 | 0.265 |
| 90 | TBG-N5 | No zone |  |  |  |
| 91 | TBG-N6 | 3 | 0.9 | 3.30 | 0.436 |
| 92 | TBG-N7 | 1.5 | 0.6 | 2.50 | 0.300 |
| 93 | TBG-N8 | 2.8 | 0.8 | 3.50 | 0.300 |
| 94 | TBG-NMP6 | No zone |  |  |  |
| 95 | TBG-NMP5 | No zone |  |  |  |
| 96 | TBG-NMP7 | 2.7 | 0.8 | 3.30 | 0.265 |
| 97 | TBG-AM1A | 1.7 | 0.6 | 2.80 | 0.100 |
| 98 | TBG-AM5 | 3.5 | 0.9 | 3.90 | 0.100 |
| 99 | TBG-AM31 | 2 | 0.8 | 2.50 | 0.265 |
| 100 | TBG-AM21 | No zone |  |  |  |
| 101 | TBG-AM27 | No zone |  |  |  |
| 102 | TBG201 | 1.7 | 0.7 | 2.40 | 0.100 |
| 103 | TBG19 | 3.2 | 0.9 | 3.50 | 0.500 |
| 104 | TBG3-6 | No zone |  |  |  |
| 105 | TBG3-8 | No zone |  |  |  |
| 106 | TBG3-12 | No zone |  |  |  |
| 107 | TBG-AM47 | No zone |  |  |  |
| 108 | TBG-AM83 | 1.3 | 0.3 | 4.30 | 0.265 |
| 109 | TBG-A6-2 | 2.4 | 0.7 | 3.33 | 0.416 |
| 110 | TBG-S14A2 | No zone |  |  |  |
| 111 | TBG-S40A5 | 3 | 1 | 3.00 | 0.200 |
| 112 | TBG-S13A5 | No zone |  |  |  |
| 113 | TBG-AG21 | No zone |  |  |  |
| 114 | TBG-AG20 | No zone |  |  |  |
| 115 | TBG-AG28 | 2 | 0.8 | 2.50 | 0.300 |
| 116 | TBG-B1 | No zone |  |  |  |
| 117 | TBG-B2 | 2.3 | 1.3 | 1.80 | 0.300 |
| 118 | TBG-B3 | No zone |  |  |  |
| 119 | TBG-B4 | No zone |  |  |  |
| 120 | TBG-B5 | No zone |  |  |  |
| 121 | TBG-WY1 | No zone |  |  |  |
| 122 | TBG-WY2 | 3.7 | 1.1 | 3.40 | 0.173 |
| 123 | TBG-WY5 | 3.3 | 0.9 | 3.70 | 0.200 |
| 124 | TBG-WY8 | 2.1 | 0.5 | 4.20 | 0.361 |
| 125 | TBG-WY9 | 3.2 | 0.9 | 3.60 | 0.265 |
| 126 | TBG-WY17 | No zone |  |  |  |
| 127 | TBG-WY19 | 2.7 | 0.7 | 3.90 | 0.265 |

\*strains selected for secondary screening

### Secondary Screening of Exo-β-1,4-Glucanase and Endo-β-1,3-Glucanase Activities

Potential isolates selected by primary screening of both exo-β-1,4-glucanase and endo-β-1,3-glucanase enzymes were subcultured and prepared spore inoculum. Secondary or quantitative enzyme screening was performed by submerged fermentation. 14 strains with exo-β-1,4-glucanase and 8 strains with endo-β-1,3-glucanase activities were selected for quantitative enzyme screening. Prepared 3×10^8^ spores.mL^-1^ were inoculated into liquid media for both enzymes. After 5 days of incubation at 28 □ in the particular screening media, the actinomycetes strain TBG-MR17 showed highest exo-β-1,4-glucanase activity and TBG-AL13 produced highest endo-β-1,3-glucanase activity. The exo-β-1,4-glucanase and endo-β-1,3-glucanase activities obtained in secondary screening are shown in table 4. The strains produced highest enzyme activities, TBG-MR17 for exo-β-1,4-glucanase activity (141 U.mL-1) and TBG-AL13 for endo-β-1,3-glucanase activity (892 U.mL-1), where considered as potent beta glucanase enzyme producers.

**Table 4.**
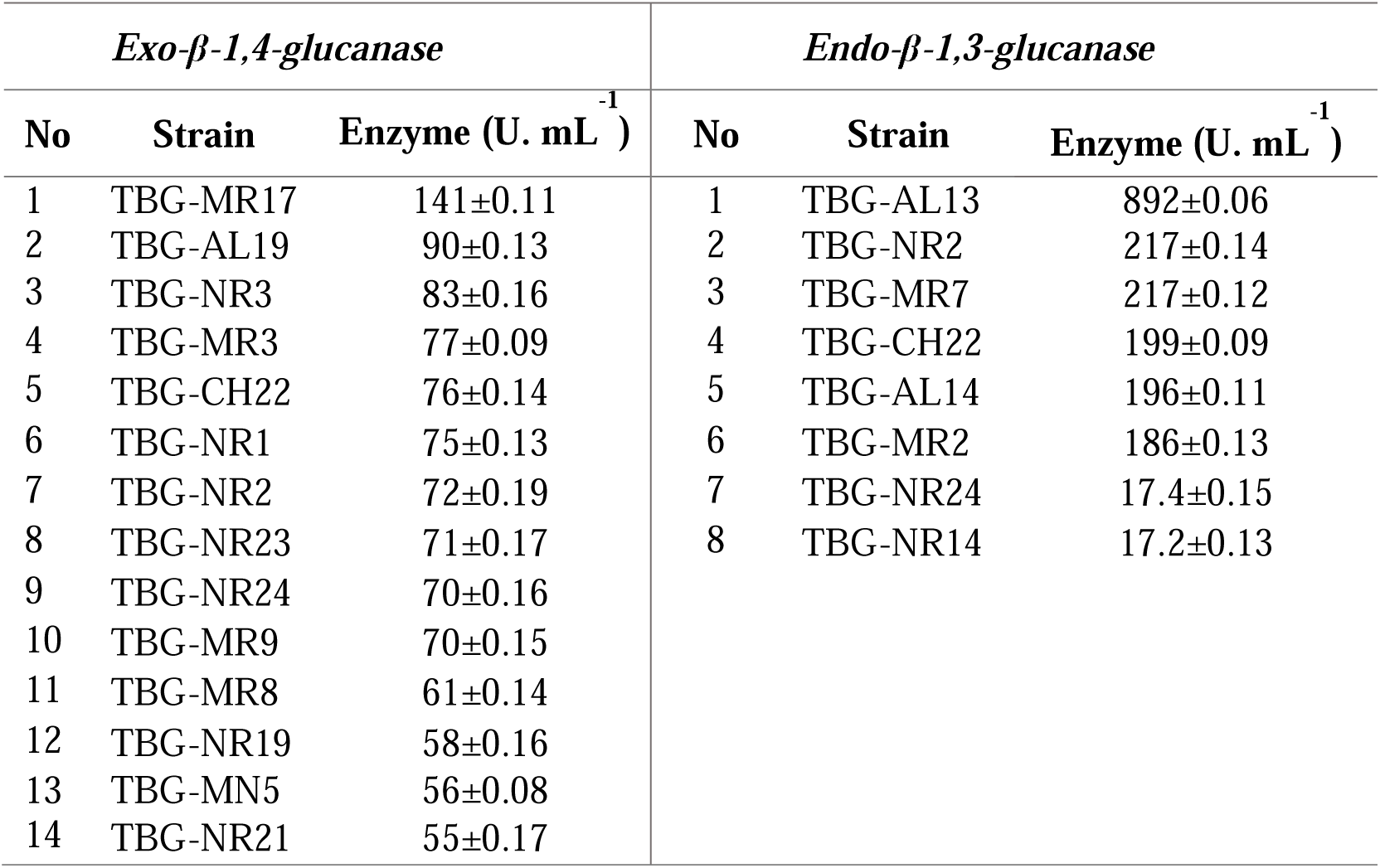
Secondary screening of enzyme activities

| *Exo-β-1,4-glucanase* |  |  | *Endo-β-1,3-glucanase* |  |  |
| --- | --- | --- | --- | --- | --- |
| No | Strain | Enzyme (U. mL<sup>-1</sup>) | No | Strain | Enzyme (U. mL<sup>-1</sup>) |
| 1 | TBG-MR17 | 141±0.11 | 1 | TBG-AL13 | 892±0.06 |
| 2 | TBG-AL19 | 90±0.13 | 2 | TBG-NR2 | 217±0.14 |
| 3 | TBG-NR3 | 83±0.16 | 3 | TBG-MR7 | 217±0.12 |
| 4 | TBG-MR3 | 77±0.09 | 4 | TBG-CH22 | 199±0.09 |
| 5 | TBG-CH22 | 76±0.14 | 5 | TBG-AL14 | 196±0.11 |
| 6 | TBG-NR1 | 75±0.13 | 6 | TBG-MR2 | 186±0.13 |
| 7 | TBG-NR2 | 72±0.19 | 7 | TBG-NR24 | 17.4±0.15 |
| 8 | TBG-NR23 | 71±0.17 | 8 | TBG-NR14 | 17.2±0.13 |
| 9 | TBG-NR24 | 70±0.16 |  |  |  |
| 10 | TBG-MR9 | 70±0.15 |  |  |  |
| 11 | TBG-MR8 | 61±0.14 |  |  |  |
| 12 | TBG-NR19 | 58±0.16 |  |  |  |
| 13 | TBG-MN5 | 56±0.08 |  |  |  |
| 14 | TBG-NR21 | 55±0.17 |  |  |  |

## Discussion

Western Ghats are one of the treasurable natural resources of earth (Gadgil, 1979). Mostly it has protected and conserved unique biodiversity, point out the chance to discover unidentified microorganisms. Study on actinobacterial diversity for β-glucanase enzymes from 10 different less explored unusual and unique ecological niche in Western Ghats regions, has led to the isolation of 127 morphologically different actinomycetes strains. We have selected unexploited Western Ghats regions in Kerala mainly natural mountain and forest areas of Wayanad, Nelliyampathy, Neriyamangalam, Munnar, Chinnar, Anamalai, Marayoor, Kulathupuzha, Palode and Agasthyarkoodam. According to Nampoothiri et al., (2013), Western Ghats soil samples were immensely utilized for the isolation of industrially important novel enzyme producing microorganisms. Forest soils are the huge domicile of taxonomically diverse actinomycetes strains especially Streptomyces sp. Despite the presence of rich recalcitrant biopolymers they are actively involved forest nutrient turnover (Bontemps et al., 2013).

Pre-treatment of soil samples with CaCO_3_ (1%) produced better colony counts. CaCO3 accelerates the growth of actinomycetes spores in the collected soils. It is one of the good method for enriching actinomycetes propagules, which significantly yielded high total plate counting. This method was also proved effective in increasing the number of rare isolates. It yielded two-fold higher number of microbes than from untreated soil samples (Tiwari and Gupta, 2012). According to Otoguro et al. (2001), calcium carbonate soil treatment yielded good colony count (2.9×105 CFU.g-1) of dried soil and leaf litter samples. CaCO_3_ alter the pH of soil, thus favours the growth of spores and stimulate the formation of aerial mycelia (Natsume et al., 1989).

Current study also intended to screen and quantify two different β-glucanase, exo-β-1,4-glucanase and endo-β-1,3-glucanase, from isolated actinomycetes strains. Avicel (microcrystalline cellulose) was used as the substrate for exo-β-1,4-glucanase. As per earlier reports, Avicel is the specific substrate for exo-acting β-1,4-glucanase (Florencio et al., 2012; Annamalai et al., 2016b). Exo-glucanase or avicelase produced by Streptomyces reticuli was efficiently utilize Avicel (crystalline cellulose), when provided Avicel as a sole carbon source (Wachinger et al., 1989; Walter and Schrempf, 1996).

Primary screening is based on the clear halo formation around the isolated colonies which directly indicated the region of enzyme action to produce glucose units. The congo red dye remain attached to areas where the presence of β-1,4-D-glucanohydrolase bonds (Lamb and Loy, 2005). Out of total 127 isolates, 106 strains (83%) produced exo-β-1,4-glucanase enzyme activity, indicated majority of Western Ghats actinomycetes isolates are good producers of exo-β-1,4-glucanase. AZCL-Pachyman was used as substrate for screening endo-β-1,3-glucanase activity. According to previous reports, pachyman is a purely β-1,3-linked substrate so can be used for determining precise endo-β-1,3-glucanase activity (Zantinge et al., 2002; Sakamoto et al., 2006). Azurine cross-linked (AZCL) polysaccharide substrates are widely used for screening glycosyl hydrolases (Li et al., 2011; Nyyssonen et al., 2013). Out of total 127 isolated strains, only 79 strains (62%) produced blue halo by the action of the endo-β-1,3-glucanase which degraded dye linked polysaccharides to monosaccharide and subsequently released dye.

Enzyme degradation in agar media was calculated in terms of enzymatic index. It is modest and fastest methodology to screen the strains prospective for enzyme production within the same genus (Ruegger and Tauk-Tornisielo, 2004). Ten et al. (2004), indicated that cellulase, xylanase and amylase producing strains effectively selected using halo zone diameter and colony diameter based enzymatic index. Strains showed high EI values (≥ 4.5) were considered as potential enzyme producers and selected for quantitative enzyme assays. Bhaturiwala et al., (2017), reported enzyme activity profiling of 20 actinomycetes strains such as cellulase, lipase, chitinase, β-mannanase, amylase, caseinase, caffeinase and xylanase activities by determining enzymatic index. Highest EI values were observed in TBG-NR3 (7.27±0.225) and TBG-CH22 (8.53±0.058) respectively for exo-β-1,4-glucanase and endo-β-1,3-glucanase plate assays.

Based on EI values 14 strains with exo-β-1,4-glucanase activity and 8 strains with endo-β-1,3-glucanase activity were selected for quantitative evaluation using submerged fermentation. After five days of incubation, actinomycetes strain TBG-MR17 showed highest exo-β-1,4-glucanase activity (95 U.mL-1) and TBG-AL13 showed the highest endo-β-1,3-glucanase activity (219 U.mL-1). Florencio et al. (2012), reported that no straight correlation was obtained between quantitative enzyme activity and enzymatic indexes. The same results were observed in our study. However quantitative method considered as final validation, as it deliver more exact data with slight variability, greater sensitivity and allow to compare relative amount of enzymes (Bisswanger, 2014; Farris et al., 2016).

## Conclusion

Microbial communities from diverse Western Ghats ecological niches are almost unexplored and rich reservoirs of valuable metabolites, likely to provide extensive applications beneficial to humanity. Especially fast growing prerequisites for enzymes in diverse extents demands an urgent need to explore actinomycetes as a treasured source of marketable enzymes. Our exploration revealed that Western Ghats ecosystems are unusual habitat for promising actinobacterial diversity with extraordinary β-glucanase activity. Extensive range of climatic environments, rich woodland areas and less discrepancies in soil type with less acidic to alkaline pH and low EC, are relatively favourable for the largest distribution of actinobacteria with high β-glucanase activity. Total of 127 actinomycetes isolates were documented during the course of study. Qualitative enzyme assay revealed, 106 strains (83%) showed exo-β-1,4-glucanase enzyme activity and only 79 strains (62%) produced endo-β-1,3-glucanase activity. According to quantitative activity profiling, the actinomycetes strains TBG-MR17 and TBG-AL13 recognised as dominant exo-β-1,4-glucanase and endo-β-1,3-glucanase producers with 141 and 892 U.mL^-1^ of respective activities. This is the first report of exploration of Western Ghats actinomycetes for both exo-β-1,4-glucanase and endo-β-1,3-glucanase enzymes with tremendously high activities.

## References

Balachandran, C., Duraipandiyan, V., & Ignacimuthu, S. (2012). Cytotoxic (A549) and antimicrobial effects of Methylobacterium sp. isolate (ERI-135) from Nilgiris forest soil, India. Asian Pacific Journal of Tropical Biomedicine, 2, 712–716.

Bhaturiwala, R. A., Jha, S. C., Jain, N. K., & Modi, H. A. (2017). Enzyme profiling of selected chitinase producing Actinomycetes. European Journal of Biotechnology and Bioscience, 5(1), 39–43.

Bisswanger, H. (2014). Enzyme assays. Perspectives in Science, 1(1-6), 41–55.

Bontemps, C., Toussaint, M., Revol, P. V., Hotel, L., Jeanbille, M., Uroz, S., Turpault, M. P., Blaudez, D., & Leblond P. (2013). Taxonomic and functional diversity of Streptomyces in a forest soil. FEMS Microbiology Letters, 342(2), 157–167.

El-Nakeeb, M. A., & Lechevalier, H. A. (1963). Selective isolation of aerobic actinomycetes. Applied Microbiology, 11(2), 75–77.

Farris, M. H., Ford, K. A., & Doyle, R. C. (2016). Qualitative and quantitative assays for detection and characterization of protein antimicrobials. Journal of Visualized Experiments, 10(110), e53819.

Fayad, K. P., Simao-Beaunoir, A. M., Gauthier, A., Leclerc, C., Mamady, H., Beaulieu, C., & Brzezinski, R. (2001). Purification and properties of a beta-1,6-glucanase from Streptomyces sp. EF-14, an actinomycete antagonistic to Phytophthora spp. Applied Microbiology and Biotechnology, 57(1-2), 117–23.

Florencio, C., Couri, S., & Farinas, C. S. (2012). Correlation between agar plate screening and solid-state fermentation for the prediction of cellulase production by Trichoderma Strains. Enzyme research, 793708, 1–7.

Fulop, L., & Ponyi, T. (1997). Rapid screening for endo-β-1,4-glucanase and endo-β-1,4-mannanase activities and specific measurement using soluble dye-labelled substrates. Journal of Microbiological Methods, 29(1), 15–21.

Gadgil, M. (1979). Hills, dams and forests. Some field observations from the Western Ghats. Proceedings of the Indian Academy of Science, 2(3), 291–303.

Hong, T. Y., Hsiao, Y. Y., Meng, M., & Li, T. T. (2008). The 1.5 A structure of endo-1,3-beta-glucanase from Streptomyces sioyaensis: evolution of the active-site structure for 1,3-beta-glucan-binding specificity and hydrolysis. Acta Crystallographica D Biological Crystallography, 64(9), 964–70.

Jalaja, V., Swetha, S., Kumar, S. S., Deepthi, A., Kumar, R. S., & Pandey, A. (2011). Isolation and characterization of alpha amylase from a metagenomic library of Western Ghats of Kerala, India. Biologia, 66, 939–944.

Lamb, J., & Loy, T. (2005). Seeing red: the use of Congo red dye to identify cooked and damaged starch grains in archaeological residues. Journal of Archaeological Science, 32(10), 1433–1440.

Li, L. L., Taghavi, S., McCorkle, S. M., Zhang, Y. B., Blewitt, M. G., Brunecky, R., Adney, W. S., Himmel, M. E., Brumm, P., Drinkwater, C., Mead, D. A., Tringe, S. G., … Lelie, D. V. (2011). Bioprospecting metagenomics of decaying wood: mining for new glycoside hydrolases. Biotechnology for biofuels, 4(1), 23.

Meddeb-Mouelhi, F., Moisana, F. J., & Beauregard, M. (2014). A comparison of plate assay methods for detecting extracellular cellulase and xylanase activity. Enzyme and Microbial Technology, 66, 16–19.

Miller, G.L. (1959). Use of dinitrosalicylic acid reagent for determination of reducing sugar. Analytical Chemistry, 31, 426–428.

Mohandas, S. P., Ravikumar, S., Menachery, S. J., Suseelan, G., Narayanan, S.S., Nandanwar, H., & Nampoothiri, K.M. (2012). Bioactives of microbes isolated from Western Ghats belt of Kerala show lactamase inhibition along with wide spectrum antimicrobial activity. Applied Biochemistry and Biotechnology, 167 (6), 1753–1762.

Nampoothiri, K. M., Ramkumar, B., & Pandey, A. (2013). Western Ghats of India: Rich source of microbial diversity. Journal of Scientific and Industrial Research, 72, 617–623.

Natsume, M., Yasui, K., & Marumo, S. (1989). Calcium ion regulates aerial mycelium formation in actinomycetes. The Journal of Antibiotics, 42, 440–447.

Nyyssonen, M., Tran, H. M., Karaoz, U., Weihe, C., Hadi, M. Z., Martiny, J. B., Martiny, A. C., …… Brodie, E. L. (2013). Coupled high-throughput functional screening and next generation sequencing for identification of plant polymer decomposing enzymes in metagenomic libraries. Frontiers in Microbiology, 4, 282.

Otoguro, M., Hayakawa, M., Yamazaki, T., & Iimura, Y. (2001). An integrated method for the enrichment and selective isolation of Actinokineospora spp. in soil and plant litter. Journal of applied microbiology, 91(1), 118–130.

Ruegger, M. J. S., & Tauk-Tornisielo, S. M. (2004). Cellulase activity of fungi isolated from soil of the ecological station of Jureia-Itatins, Sao Paulo, Brazil. Brazilian Journal Botany, 27, 205–211.

Sakamoto, Y., Watanabe, H., Nagai, M., Nakade, K., Takahashi, M., & Sato, T. (2006). Lentinula edodes tlg1 encodes a thaumatin-like protein that is involved in lentinan degradation and fruiting body senescence. Plant Physiology, 141(2), 793–801.

Sakdapetsiri, C., Fukuta, Y., Aramsirirujiwet, Y., Shirasaka, N., & Kitpreechavanich, V. (2016). Antagonistic activity of endo-β-1,3-glucanase from a novel isolate, Streptomyces sp. 9×166, against black rot in orchids. Journal of Basic Microbiology, 56(5), 469–79.

Shi, P., Yao, G., Yang, P., Li, N., Luo, H., Bai, Y., Wang, Y., & Yao, B. (2010). Cloning, characterization, and antifungal activity of an endo-1,3-beta-D: -glucanase from Streptomyces sp. S27. Applied Microbiology and Biotechnology, 85 (5), 1483–1490.

Ten, L. N., Im, W. T., Kim, M. K., Kang, M. S., & Lee, S. T. (2004). Development of a plate technique for screening of polysaccharide-degrading microorganisms by using a mixture of insoluble chromogenic substrates. Journal of Microbiological Methods, 56(3), 375–382.

Tiwari, K., & Gupta, R. K. (2012). Diversity and isolation of rare actinomycetes: an overview. Critical Reviews in Microbiology, 39(3), 256–294.

Velho-Pereira, S. & Kamat, N. M. (2013). Actinobacterial research in India. Indian Journal of Experimental Biology, 51, 573–596.

Wachinger, G., Bronnenmeier, K., Staudenbauer, W. L., & Schrempf, H. (1989). Identification of mycelium-associated cellulase from Streptomyces reticuli. Applied Environmental Microbiology, 55, 2653–2657.

Walter, S., & Schrempf, H., (1996). Physiological studies of cellulase (avicelase) synthesis in Streptomyces reticuli. Applied Environmental Microbiology, 62(3), 1065–1069.

Woo, J. B., Kang, H. N., Woo, E. J., & Lee, S. B. (2014). Molecular cloning and functional characterization of an endo-β-1,3-glucanase from Streptomyces matensis ATCC 23935. Food Chemistry, 1(148), 184–187.

Wu, H., Shimoi, H., & Ito, K. (2002). Purification and characterization of beta-1,6-glucanase of Streptomyces rochei application in the study of yeast cell wall proteins. Bioscience Biotechnology and Biochemistry, 66(11), 2515–2519.

Wu, Q., Dou, X., Wang, Q., Guan, Z., Cai, Y., & Liao, X. (2018). Isolation of β-1,3-Glucanase-Producing Microorganisms from Poria cocos Cultivation Soil via Molecular Biology. Molecules, 23(7), 1555.

Zantinge, J. L., Huang, H. C., & Cheng, K. J. (2002). Microplate diffusion assay for screening of beta-glucanase-producing microorganisms. Biotechniques, 33(4), 798–806.

